# Systems analysis reveals neuregulin-1 control of cardiomyocyte size and shape mediated by distinct PI3K and p38 pathways

**DOI:** 10.1101/2025.10.01.679873

**Authors:** Pichayathida Luanpaisanon, Karen A. Ryall, Philip M. Tan, James O’Hearn, Laura A. Woo, Bryana N. Harris, Bethany Wissmann, Alexander Paap, Matthew Rhoads, Mohammad Fallahi-Sichani, Jeffrey J. Saucerman

## Abstract

Pathological and physiological stresses induce diverse forms of cardiac hypertrophy, with distinct manifestations in cardiomyocyte size and shape regulated by still poorly understood signaling networks. Here, we combined high-content morphological profiling, phospho-protein arrays, and systems modeling to characterize the diverse forms of hypertrophy induced by angiotensin II, endothelin-1, insulin growth factor-1, and neuregulin-1. Reverse-phase protein array profiling of neonatal rat cardiomyocytes and partial least squares regression modeling revealed that Akt, GSK3, and MAPK signaling are differentially regulated by hypertrophic agonists and are predictive of distinct phenotypic outcomes. Among these agonists, neuregulin-1 uniquely induced cardiomyocyte elongation in both neonatal rat and human iPSC-derived cardiomyocytes, in addition to increasing cell area. Pharmacological perturbations in neonatal rat cardiomyocytes demonstrated that neuregulin1-induced elongation and area expansion both require PI3K activity, whereas p38 selectively mediates cell area. A logic-based network model incorporating dual-specificity phosphatases were sufficient to capture the amplifying PI3K and transient p38 signaling dynamics driving phenotypic changes. Together, these results identify distinct signaling cascades by which neuregulin-1 coordinates cardiomyocyte size and shape, providing mechanistic insight into how hypertrophic remodeling can be differentially regulated. This systems approach provides new insight into the pathways that drive distinct forms of cardiomyocyte hypertrophy, highlighting opportunities to selectively target maladaptive remodeling in heart failure.

**Highlights:** -Reverse-phase protein arrays capture distinct signatures of cellular signaling in response to diverse hypertrophic ligands.

-Partial least squares regression model maps proteomic signatures to diverse patterns of cell morphology and gene expression.

-Combinatorial ligand-inhibitor screen validates predicted causal regulators of mRNAs and cell morphology

-PI3K mediates both neuregulin-1-induced elongation and cell area, validating the PLSR model. In contrast, p38 regulates cell area but not elongation.

-Logic-based model demonstrates that the characterized mechanisms are sufficient to predict how distinct PI3K and p38 dynamics drive size and shape.

## Introduction

Cardiac hypertrophy is a compensatory response of the heart to increased workload, characterized by an increase in cardiomyocyte size and increased cardiac mass. Cardiomyocytes account for approximately 30% of total cell number but contribute to 70–80% of the heart’s mass [1], [2]. Hypertrophy can be a response to either pathological stresses (e.g. myocardial infarction, hypertension) or physiological stresses (e.g. exercise, pregnancy), in which cardiomyocytes undergo considerable changes in cell size and shape [2]. While the signaling pathways driving cardiomyocyte size have been studied intensely, the mechanisms governing changes in cardiomyocyte shape remain poorly understood. For example, pathologic hypertrophy such as hypertension causes a pressure overload that increases cross-sectional area, resulting in concentric hypertrophy [3], [4]. Other pathologic stimuli such mitral regurgitation cause volume overload, which elicits eccentric hypertrophy through a predominate lengthening of individual cardiomyocytes, which is key to the transition to heart failure [5], [6]. At the same time, physiological cues such as resistance and endurance exercise induce either concentric or eccentric hypertrophy, driven by molecular pathways that are less characterized [7], [8].

Cardiomyocyte hypertrophy is sensitive to both mechanical and chemical stimuli, including cytokines, growth factors, catecholamines, vasoactive peptides, and hormones. Among the most well-characterized hypertrophic growth factors are angiotensin II (AngII), endothelin-1 (ET1), neuregulin-1 (Nrg1), and insulin growth factor-1 (IGF1) [9], [10], [11], [12], [13]. Phenotypic screens comparing across stimuli have revealed ligand-specific regulation of cardiomyocyte size, shape, and other properties. Bass et al. performed an early developed early high-content image analyses that revealed that of 5 stimuli that increased cardiomyocyte area, only phenylephrine and serum increased sarcomere organization associated with physiological hypertrophy [14]. In a subsequent screen of 15 hypertrophic agonists, Ryall et al. found that Nrg1 induced the greatest cardiomyocyte elongation and CITED4 mRNA. Knockdown of CITED4 further accentuated elongation, revealing an incoherent feedforward loop [13]. However, in most cases it is still unclear how interconnected signaling pathways induce distinct cell features associated with hypertrophy, especially for cell shape.

To identify pathways that distinctly regulate cardiomyocyte size, shape, and gene expression, we performed a proteomic screen under AngII, ET1, Nrg1, and IGF1 conditions where we previously found diverse cardiomyocyte phenotypes. Proteomic analysis revealed distinct cell states depending on treatment and timepoint. We further mapped the phospho-protein array data to the previous phenotypic screen to infer signaling modules and pathways. Finally, we validated the predicted mediators of Nrg1-dependent cardiomyocyte size and shape using rodent and human iPSC-derived cardiomyocytes.

## Results

### Hypertrophic ligands induce diverse phosphoprotein signatures predictive of cardiomyocyte morphology and gene expression

Previously, we performed a phenotypic screen that revealed hypertrophic ligands that induce diverse forms of cardiomyocyte morphology and gene expression [13]. That study found a number of new ligand-specific relationships such as an incoherent feedforward with Nrg1- and CITED4-regulated myocyte elongation regulated by CITED4, as well as CTGF regulating AngII-induced Bax expression [13]. However, the signaling pathways driving these responses were not characterized. To identify signatures of protein phosphorylation and abundance that may drive these phenotypic responses, we treated neonatal rat cardiomyocytes with Nrg1 (10 ng/mL), AngII (1 µM), ET1 (100 nM), or IGF (10 nM) as previously, but here measuring 172 total and phosphor- protein antibody probes by reverse phase protein array (RPPA) at 1 h and 48 h. Heatmaps summarizing this data revealed both time- and ligand-specific proteomic responses compared to serum-free controls **(Figure 1A)**. To highlight dynamic changes that are unique among treatment groups, from each group we selected the top 20% proteins most expressed at 1 hr and 48 hr.

**Figure 1.**
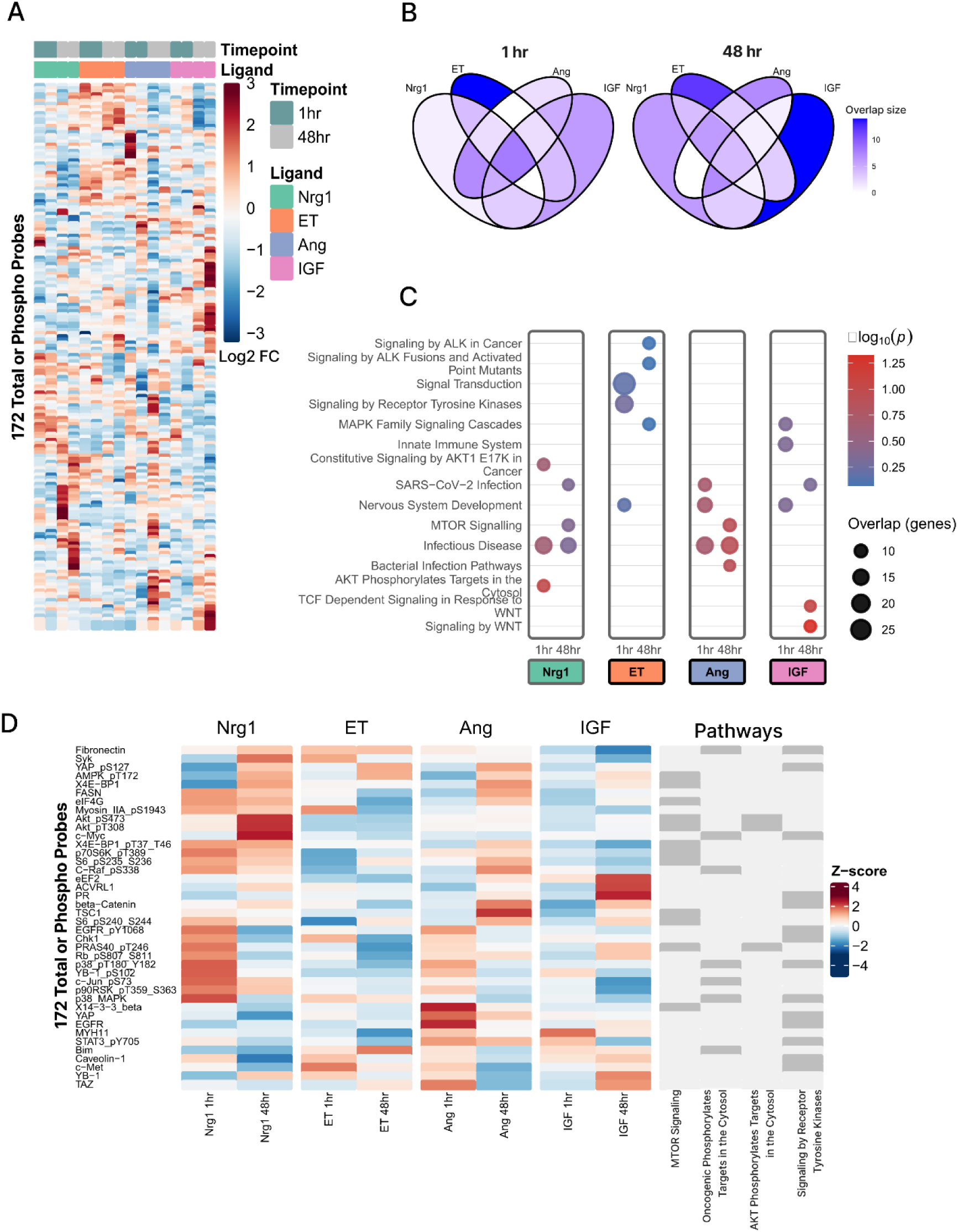
Hypertrophic ligands induce diverse changes in protein abundance and phosphorylation in neonatal rat cardiomyocytes. (A) Hierarchical clustering of RPPA data from 172 total/phospho probes following 1 hr and 48 hr treatment with Nrg1 (10 ng/mL, R&D #396-HB), Ang II (1 µM), ET1 (100 nM), or IGF1 (10 nM) (log2 fold-change compared to control). For each timepoint, data were collected from 2 independent isolations and were subsequently averaged. (B) Venn diagrams of top 20% most responsive proteins under each treatment at 1 hr vs 48 hr. (C) Enrichr Reactome pathway enrichment analysis, showing the top 3 enriched pathways with overlap genes ≥ 10. (D) Differentially expressed or phosphorylated proteins, annotated by Reactome pathways most enriched for each treatment.

Venn diagrams revealed that many responses were largely shared across ligands at 1 hr, but proteomic responses became progressively more ligand-specific by 48 hr **Figure 21)**. For example, Nrg1 and AngII shared 22 differentially-expressed proteins at 1 hr and none at 48 hr. ET was the only condition with a substantial set of unique proteins already at 1 hr. Pathway enrichment using Enrichr (Reactome database) [16, 17] identified enrichment of differentially expressed proteins associated with AKT/mTOR signaling under Nrg1 treatment, receptor- tyrosine-kinase–related pathways and MAPK signaling for ET treatment, infection/immune- related pathways for AngII treatment, and developmental signaling and WNT signaling for IGF treatment **(Figure 1C)**. Exploration of differentially expressed proteins associated within each of these enriched pathways revealed that Nrg treatment induces early activation of the Ras/MAPK signaling—elevated phosphorylation of ERK/MAPK, p38, JNK, c-Jun, and c-Raf—at 1 hr, whereas AKT was significantly upregulated by 48 hr (**Figure 1D**).

The phospho-protein array measurements in **Figure 1** identified distinct signatures of protein abundance and phosphorylation, but these measurements alone are insufficient to determine which proteins most closely associate with cardiomyocyte morphology and gene expression. To address this gap, we mapped proteomic responses (from **Figure 1**) to our previous screen of cardiomyocyte morphology and gene expression [13] using a supervised machine learning model, partial least squares regression (PLSR). PLSR provides a powerful framework for dimensionality reduction and visualization of correlations, thereby offering insight into the organization of signaling networks [15].

To descriptively visualize the covariation structure of the combined data, we first plotted the scores for the treatment conditions together with the loadings for all proteins and the most predictive phenotype loadings in latent space (**Figure 2A**). Consistent with trends observed in the RPPA dataset alone (**Figure 1**), the treatments induced distinct protein and phenotypic across the latent space, suggesting potential ligand-specific signaling programs. Of note, Nrg1 treatment aligned strongly with cardiomyocyte elongation and CITED4 expression, consistent with our prior findings [13]. Further, unsupervised k-means clustering of protein loadings revealed a distinct module of protein expression and phosphorylation (red cluster) that are associated with cardiomyocyte elongation. Indeed, the red cluster is composed of many components of the PI3K pathway including phospho-PI3K, phospho-Akt, phospho-PDK1, phospho-GSK3, phospho-mTOR, and phospho-p70S6K. Also, within the red cluster and correlated with elongation are several MAPK superfamily proteins such as phospho-p38, phospho-MEK1, phospho-MAPK, and phospho-p90RSK.

**Figure 2.**
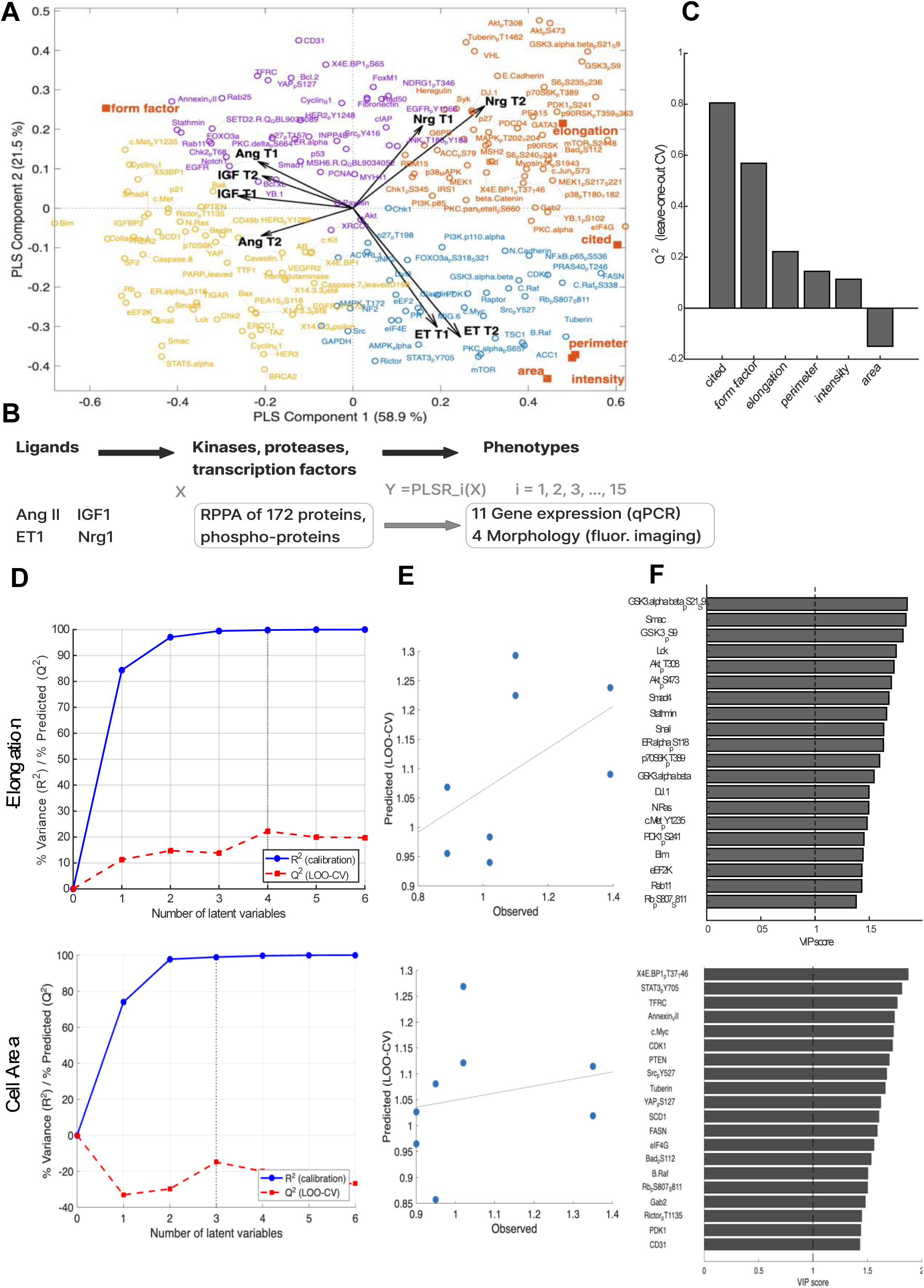
Partial least squares regression (PLSR) model predicts unique signaling pathways associated with neonatal rat cardiomyocyte size and elongation. (A) Descriptive plot of PLSR loadings for all proteins (172 predictor variables; circle markers) and the top gene expression and morphology features (6 response variables with Q^2^ ≥ 0.149). Proteins were grouped via k-means clustering (k=4). Arrows show orientation and relative magnitude of PLSR scores for ligand inputs at two timepoints (1hr and 48 h after treatment). (B) Schematic for building phenotype-specific PLSR models that use 172 protein and phospho-protein measurements (from Figure 1) to predict expression each response variable: 11 genes (by qPCR at 48 h) or 4 morphological features (by live tracking for 48 h) (both reported in [13]). (C) Leave- one-out cross validation performance (Q^2^) of the top 6 phenotype-specific PLSR models (Q^2^>0 except - 0.149 for cell area. (D) Performance of myocyte elongation and cell area PLSR models quantified by percent variance explained (R²) and leave-one-out cross-validation predictive value (Q²) as a function of latent variables. (E) Comparison of observed vs. leave-one-out predicted values for elongation and cell area using 4 and 3 latent variables, respectively. (F) Top 20 most predictive proteins ranked by VIP score for the myocyte elongation and cell area PLSR models. Dashed line indicates VIP > 1 threshold for influential predictors.

To quantify the proteins that predict each phenotype, we developed 15 phenotype-specific PLSR models using all 172 probes across four hypertrophic ligands (1 hr and 48 hr) as predictors and then each of the 11 mRNAs or four morphology metrics as the response variable (**Figure 2B**). We then performed leave-one-out cross validation to assess model performance. PLSR models for five response variables exhibited predictive ability (Q^2^>0): Cited4 mRNA, form factor, elongation, perimeter, and GFP intensity, outperforming the more commonly studied metric of cell area (Q^2^ = -0.149) (**Figure 2C**). Both elongation and cell area PLSR models explained a high level of variance in the data (R^2^) by 3 latent variables. But only the elongation model exhibited consistent leave-one-out cross-validation scores with Q^2^>0, peaking with 4 latent variables (**Figure 2D**). The higher predictive ability of the elongation PLSR model was further supported by comparing higher correlation observed vs. predicted myocyte elongation across individual experiments (**Figure 2E**).

To further dissect proteins that are most predictive of cell elongation vs. elongation, we computed VIP scores for the corresponding phenotype-specific PLSR models. Protein phosphorylation measurements most predictive of myocyte elongation were highly enriched in the PI3K/PDK1/Akt pathway, including GSKα/β(S21/S9), Akt(T308,S473), p70S6K(T389), and PDK1(S241) (**Figure 2F**). While not in the top 20, p38(T180), MEK1(S217/S22), and MAPK(T202/Y204) were also predictive of myocyte elongation (VIP scores of 1.32, 1.17, and 1.16). In contrast, protein phosphorylations most predictive of cell area were distinct, including X4E.BP1(T37/T46), Src(Y527), and YAP(S127) which have previously been associated with cardiomyocyte hypertrophy [16], [17], [18]. Taken together, these findings demonstrate that Nrg1- induced changes in phenotype are most strongly predicted by the PI3K/Akt signaling pathway and perhaps MAPK signaling modules, highlighting these pathways as potential regulators of cardiomyocyte elongation.

### MEK, PI3K, and p38 mediate distinct ligand-specific phenotypes

Given the predicted unique associations of ERK, PI3K, or p38 with elongation and other shape features (**Figure 2**), we hypothesized that inhibiting these pathways would modulate ligand-induced cardiomyocyte morphologies and gene expression. Therefore, we performed a PCR screen of 8 mRNAs that were most responsive in our previous ligand screen [13], but this time in response to 16 combinations of ligands Nrg1, ET, or AngII with inhibitors of MEK (0.1 µM PD325901), p38 (10 μM SB203580), or PI3K (10 μM LY294002) (**Figure 3A**). Broadly, mRNA abundances in response to ligands alone were consistent with the previous study [13], for example strong BNP expression and CITED4 mRNAs in response to Nrg1 and ET1 (**Figure 3B**). These responses were amplified by p38 inhibition, while on the other hand Nrg1-induced CITED4 expression was suppressed by inhibition of MEK or PI3K. These findings indicate that MEK, PI3K, and p38 may contribute to our previously characterized Nrg1-CITED4-elongation feedforward loop [13]. P38 inhibition also amplified the effects of Nrg1 on VEGF and Bcl2 (**Figure 3B**).

**Figure 3.**
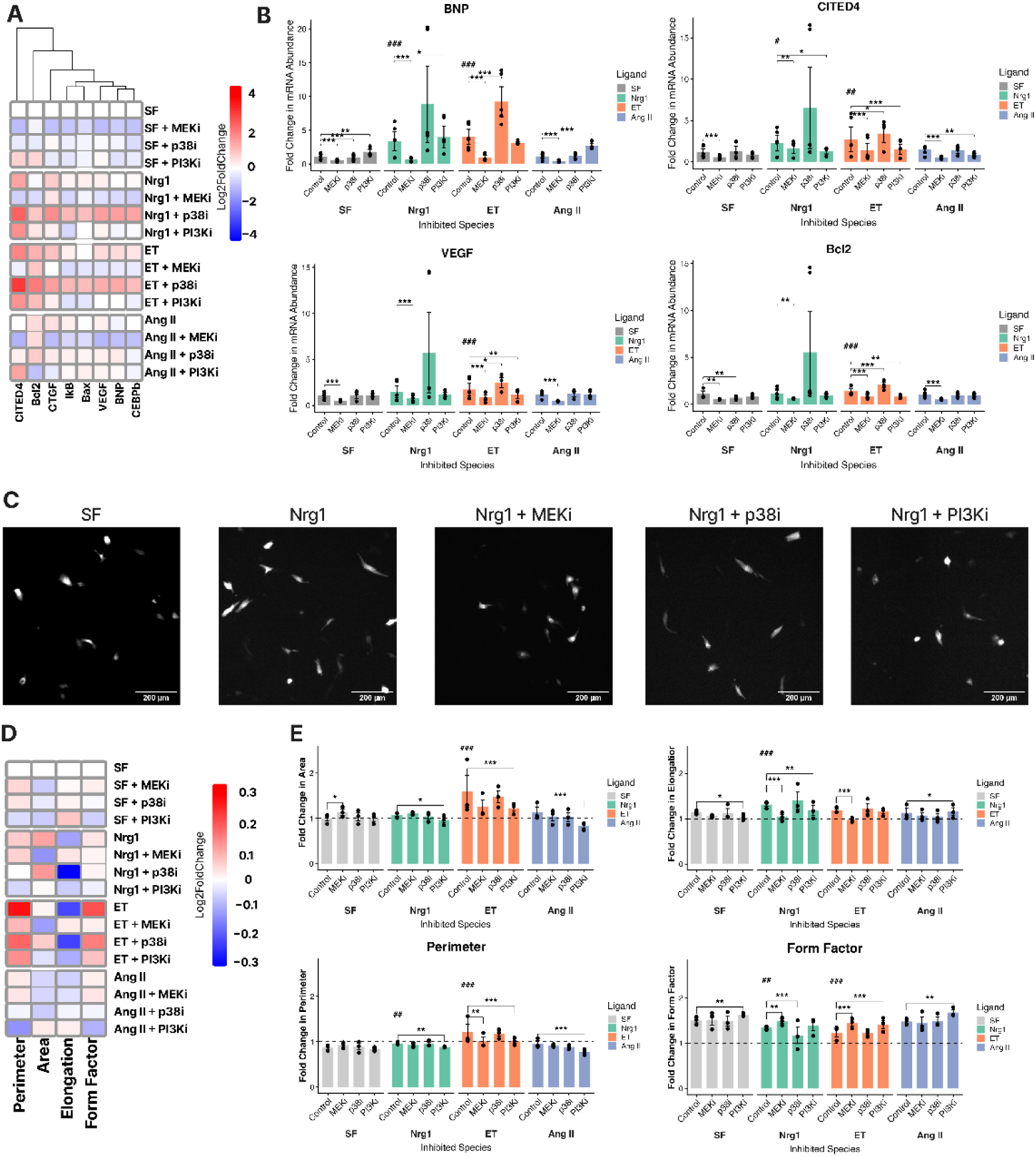
MEK, p38, and PI3K inhibition distinctly mediate ligand-induced neonatal rat cardiomyocyte gene expression and morphology. A) Heatmap showing overall mRNA responses in a ligand-inhibitor screen. Myocytes were treated for 48 h with 10 ng/mL Nrg1 (R&D #396-HB), 10 nM ET1, 1 µM AngII, or serum free negative control, in combination with inhibitors for MEK (0.1 µM PD0325901,) p38 (10 μM SB203580), or PI3K (10 μM LY294002). B) Quantitative changes in mRNA abundance for BNP, CITED4, VEGF, and Bcl2, with mRNA aggregated from 2 wells for each of 3 isolations. C) Representative images of cardiomyocytes transiently transfected with GFP under the cTnT promoter, enabling live-cell tracking of morphology. D) Heatmap showing overall morphologic responses in the ligand-inhibitor screen. E) Quantitative changes in cardiomyocyte morphology metrics, where each dot represents the median single-cell response from ∼200 myocytes for each of 3 isolations. Significance was assessed using a linear-mixed effects model (treatment group as fixed effect, isolation as random intercept). # indicates the ligand Control differs from SF Control; * indicates an inhibitor-treated condition differs from its own ligand’s Control. Significance levels for both symbols. * or # for p < 0.05, ** or ## for p < 0.01, *** or ### for p < 0.001.

We next asked whether MEK, p38, or PI3K inhibitors also modulate cardiomyocyte morphology. We again used the live-cell morphology tracking assay (representative images in **Figure 3C**), which images cells transfected with a plasmid expressing GFP under a troponin T promoter [13]. We measured cardiomyocyte area, elongation, form factor, and perimeter in response to the same 16 combinations of Nrg1/ET/AngII ligands and MEK/PI3K/p38 inhibitors. Overall, phenotypic responses to ligands without MEK/PI3K/p38 inhibitors were consistent with our previous study [13], with the strongest responses being ET1-induced cell area and perimeter as well as Nrg1-induced elongation (**Figure 3D**). In conditions of Nrg1, ET1, or AngII treatment, PI3K inhibition was most effective in reduced cell area and perimeter. In contrast, Nrg1-dependent elongation was suppressed by MEK or PI3K inhibition but enhanced by p38 inhibition (**Figure 3E**). Overall, these results validate the predicted causal roles of MEK, PI3K, and p38 anticipated from the PLSR model, and further show that correlated phospho-proteins such as PI3K and p38 have distinct causal effects on phenotype (e.g. elongation).

### PI3K mediates Nrg1-induced elongation and cell area, while p38 mediates Nrg1-induced increase in cell area

While the live-cell phenotypic screen in **Figure 3** is sensitive to single-cell changes, a limitation is that it measures only from ∼5% of GFP expressing cells which may not be representative. Therefore, to determine whether Nrg1 also induced size and shape changes in a broader population of cells, we treated rat neonatal cardiomyocytes and two commercial preparations of human iPSC-derived cardiomyocytes (iCell hiPSC-CMs and hiPSC-CM^2^ from FujiFilm) (**Figure 4A**). At 48 h, we fixed and immune-stained the cells for α-actinin, followed by high-content cell segmentation and extraction of cell morphological features. By high-content analysis, we profiled 12 single-cell size and shape features for sensitivity to Nrg1 response. Of these, the most sensitive metric to Nrg1 was the metric Feret Elongation (**Figure 4B**). The geometric measure Feret Diameter was developed by L.R. Feret in the 1930s to quantify overall extent of an object in a given direction [19]. We intentionally designed Feret Elongation to reflect the ratio of a cell’s overall extent of length (Max Feret Diameter) and overall extent of width (Min Feret Diameter), analogous to conventional manual measurements of elongation for mature cardiomyocytes (**Figure 4C**). We also showed that Feret Elongation is more sensitive in measuring elongation of idealized shapes compared to other shape measures that compare cells to circles (form factor) or ellipses (eccentricity) (**Supplementary Figure 1**). Nrg1 consistently increased Feret Elongation in this fixed-cell assay that captures all cardiomyocytes (**Figure 4D**), consistent with the live-cell assay of GFP+ transfected cardiomyocytes (**Figure 3C**). The fixed cell assay additionally showed Nrg1-dependent increase in cell area (**Figure 4E**), which was not seen in the live assay (**Figure 3C**) perhaps due to a lower sensitivity. We further tested whether Nrg1 induces elongation of human iPSC-derived cardiomyocytes, which was confirmed with two commercial products FujiFilm iCell Cardiomyocytes (hiPSC-CMs) and iCell Cardiomyocytes^2^ (hiPSC-CMs^2^) (**Supplementary Figure 2**). These results demonstrate that Nrg1-dependent cardiomyocyte elongation is a robust finding across neonatal rat and human iPSC-derived cardiomyocytes.

**Figure 4.**
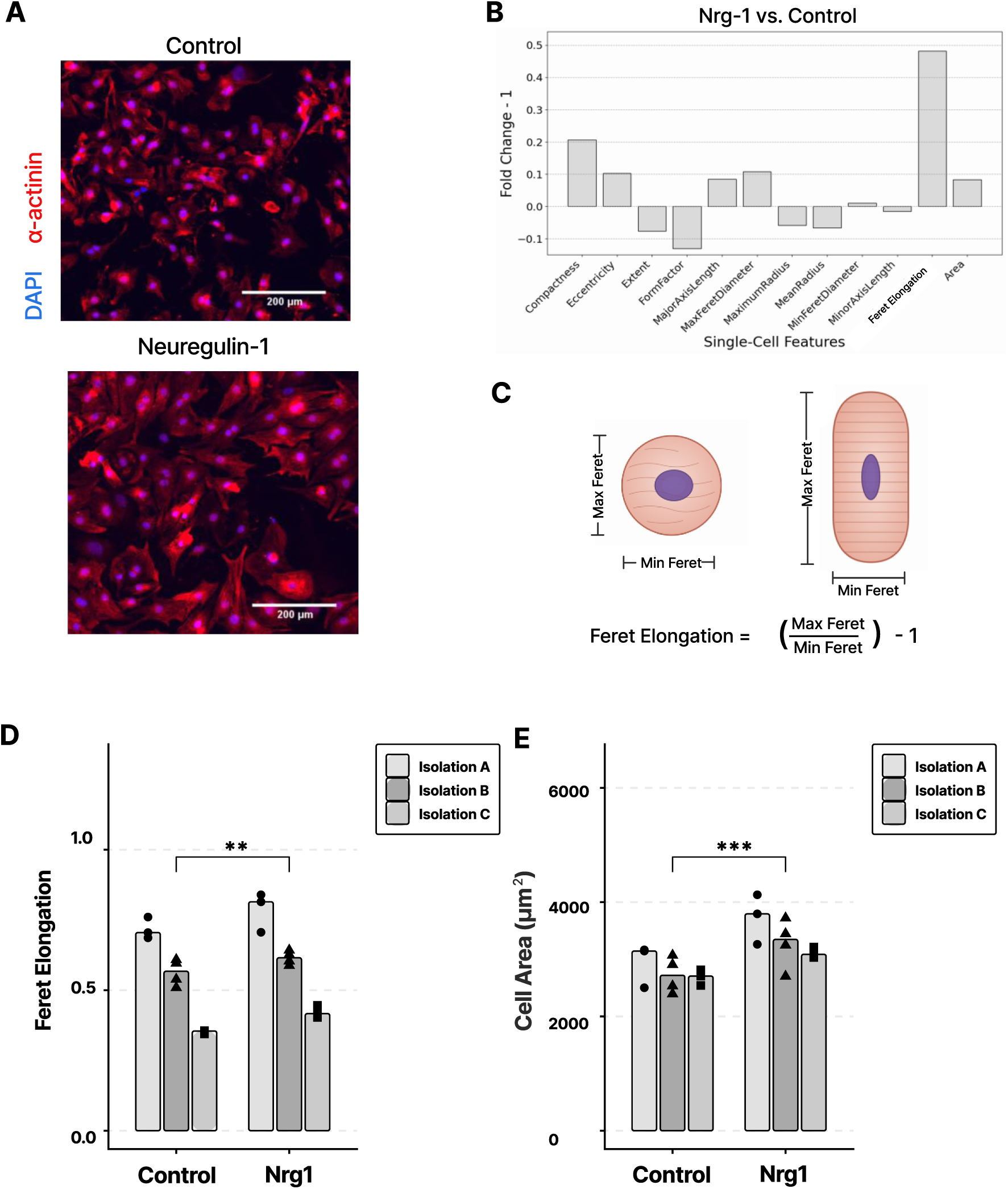
Neuregulin induces neonatal rat cardiomyocyte elongation in a more inclusive and sensitive fixed-cell assay. (A) Representative images of neonatal rat cardiomyocytes (NRCM) at 48 h after treatment with control or 100 ng/mL Neuregulin-1 (Nrg1, R&D #377-HB). Cardiomyocyte marker (α- actinin) and nuclei (DAPI) are shown. (B) Profiling 12 single-cell morphological features for sensitivity to Nrg1-treatment compared to control NRCMs, quantified as fold-change from control minus 1. (C) Schematic of how the Feret Elongation metric works for irregular geometries, using cartoons of animal ferrets for illustration only. Application to idealized circles, ellipses and rectangles are shown in Supplementary Figure 1. (D) Quantitative change in Feret Elongation in response to Nrg treatment in NRCM. (E) Quantitative change in cell area in response to Nrg treatment in NRCMs. Each point (dot, triangle, or square) reflects the mean response of ∼400 cells per well, and with bars showing mean response from across 3-4 wells/treatment from three cardiomyocyte isolations. Significance was assessed by linear-mixed effects model (treatment group as fixed effect, isolation as random intercept), with * p < 0.05, ** p < 0.01, *** p < 0.001.

Phospho-protein array measurements (**Figure 1**) and PLSR modeling (**Figure 2**) indicated that signatures of protein phosphorylation within the PI3K/AKT and MAPK superfamilies were closely aligned with Nrg-dependent cell elongation, which was modulated by MEK, PI3K, and p38 inhibitors using a live-cell phenotypic screen (**Figure 3**). We next asked whether the opposing role of PI3K and p38 inhibitors on Nrg-dependent elongation would further reproduce in the broader cell population accessible with fixed cell immunofluorescence. To test this question, we treated NRCMs with 10 ng/mL Nrg1 and applied pharmacological inhibitors for PI3K (10 μM LY294002) or p38 (10 μM SB203580) for 48 h, followed by fixation, immunofluorescence, and high-content image analysis. Consistent with our prior NRCM experiments in complementary live-cell and fixed-cell assays, Nrg1 treatment increased both cardiomyocyte elongation and cell size (**Figure 5A-C**). PI3K inhibition significantly attenuated Nrg1-induced elongation and reduced overall cell area, indicating a broad role in Nrg1-mediated hypertrophy (**Figure 5A-C**). In contrast, p38 inhibition reduced the Nrg1-dependent increase in cell area without significantly affecting elongation (**Figure 5A-C)**. Alone, PI3K inhibition decreased Feret elongation, while p38 inhibition did not. Neither inhibitor alone affected cell area.

**Figure 5.**
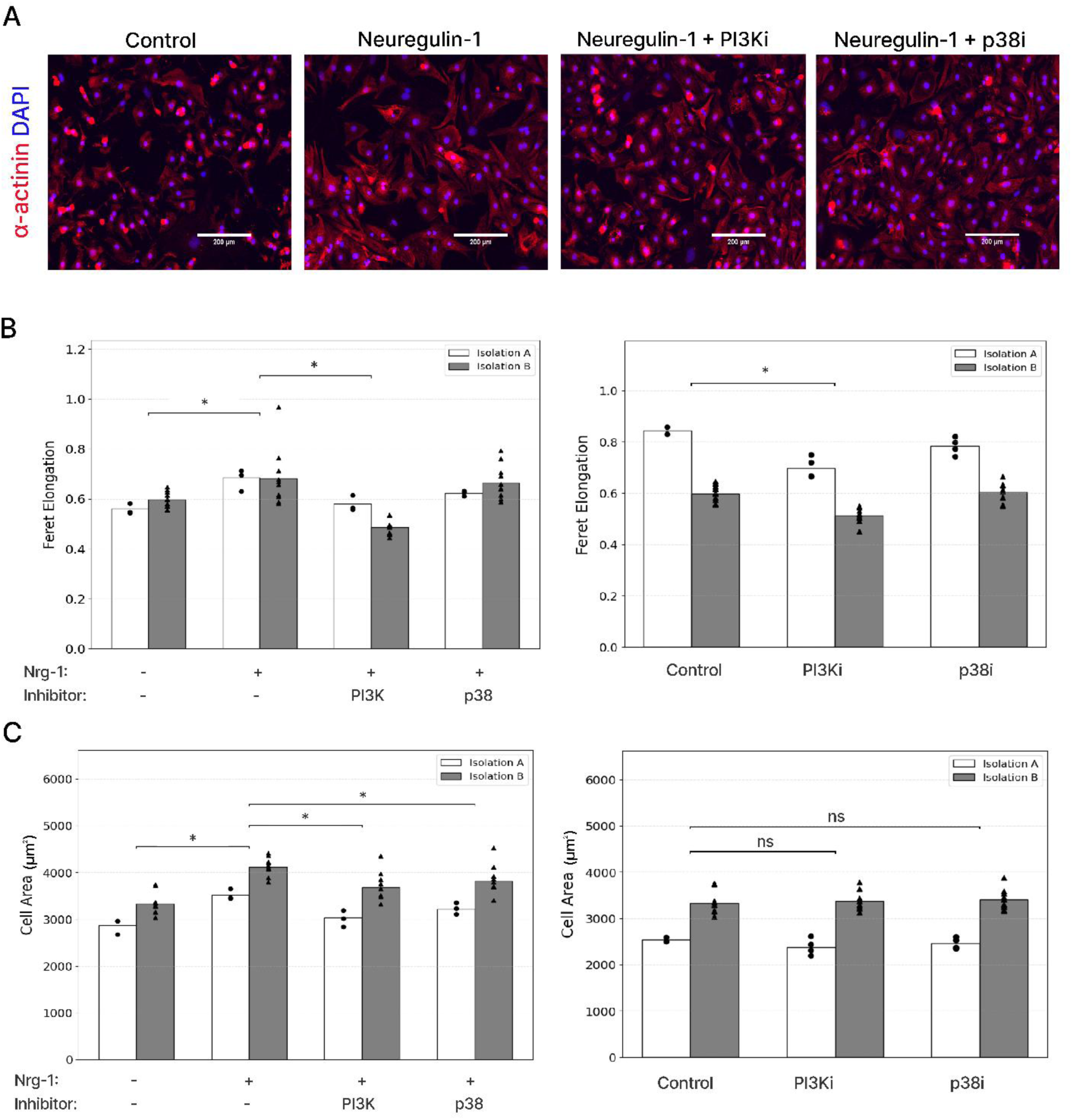
Both PI3K and p38 inhibition mitigate Nrg1-dependent cell size, but only PI3K inhibition reduces cell elongation in neonatal rat cardiomyocytes. (A) Representative images of neonatal rat cardiomyocytes (NRCMs) under four treatments: Control, 100 ng/mL Neuregulin-1 (Nrg1, R&D #377-HB), Nrg1 + 10 µM LY294002 (PI3Ki), and Nrg + 10 µM SB203580 (p38i). Cardiomyocyte marker (α-actinin) and nuclei (DAPI) are shown. (B) Quantification of Feret elongation and cell area (C) in response to PI3K and p38 inhibition, with and without Nrg1 treatment. Each point (dot, triangle, or square) reflects the mean response of ∼200 cells per well, and with bars showing mean response from across 3-4 wells/treatment from two independent cardiomyocyte isolations. Significance was assessed by linear-mixed effects model (treatment group as fixed effect, isolation as random intercept), with * p < 0.05.

### Network modeling predicts signaling cascades sufficient to regulate elongation and cell area following Nrg1 simulation

The above live-cell and fixed-cell experimental studies revealed that Nrg1 induces protein changes associated with cardiomyocyte elongation and that elongation and cell area, and that these phenotypes differentially require PI3K and p38 activity. We next asked whether our experimentally measured dynamics of PI3K and p38 are sufficient to predict the distinct network dynamics of elongation and cell area, with and without inhibitors of those pathways.

To address this question, we developed a logic-based computational model to formalize our network hypothesis, integrating data from RPPA measurements, phenotypic assays, and PLSR predictions. In this network model, Nrg1 was assumed to act through ERBB receptors, activating both PI3K/Akt/elongation/area and Ras/p38/area cascades. P38 activity was further constrained by negative regulation through Dual-specificity phosphatase (DUSP), which we hypothesized would be sufficient to predict the adaptive decline in p38 phosphorylation observed experimentally. We implemented this network structure in Netflux, a logic-based differential equation modeling platform [20].

Simulations of Nrg1 treatment predicted network-wide dynamic responses, including early activation of ERBB/Ras/p38 signaling, followed by a subsequent rise in PI3K signaling and DUSP- induced decline in p38 signaling (**Figure 6A-B**). As seen in the RPPA experiments, the model predicted that Akt phosphorylation rose slowly, while p38 phosphorylation was transient (**Figure 6C**). Perturbation simulations further validated the model. As measured experimentally, PI3K inhibition attenuated Nrg1-induced elongation and decreased cell area, while p38 inhibition selectively reduced cell area response without strongly impacting elongation **(Figure 6D).** Together, these findings demonstrate that simplified network model grounded in our experimental data recapitulates the differential pathway contributions of PI3K and p38 to both cell elongation and area.

**Figure 6.**
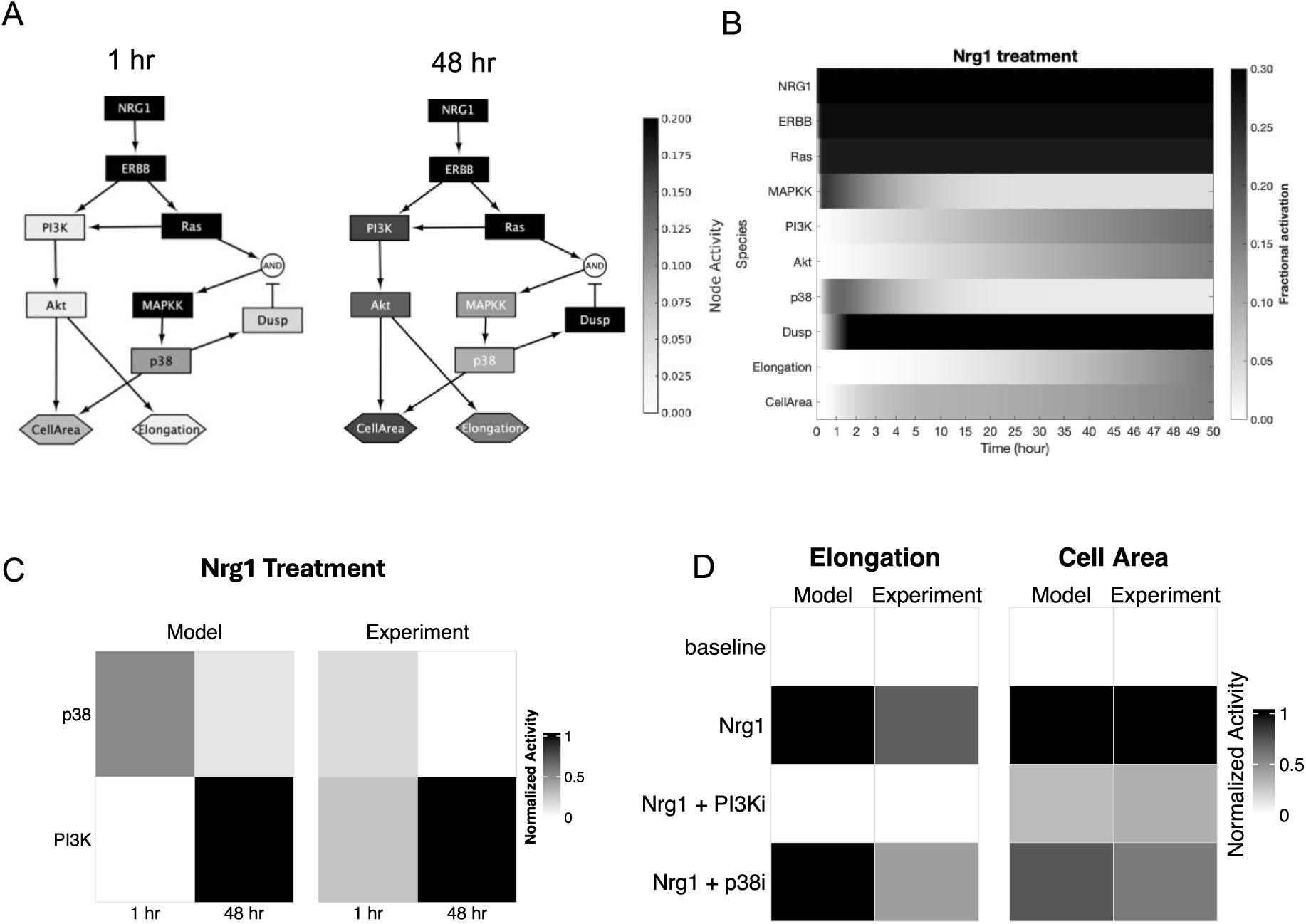
Network model identifies mechanisms sufficient for distinct regulation of cell size and shape. (A) Logic-based network model of Nrg1 signaling, with nodes colored based on their normalized activity at 1 hr and 48 hr following Nrg1 treatment. (B) Simulated response of the network in response to Nrg1. (C) Comparison of phosphorylation levels between model predictions and experiments (Akt phosphorylation: average of pS473 and pT308; p38 phosphorylation: pT180/Y182), normalized by min/max values per experiment (D) Comparison of cell area and elongation between model predictions and experimental results, normalized by min/max values per experiment.

## Discussion

A quantitative understanding of cardiomyocyte size and shape regulation is critical for elucidating the molecular pathways that govern cardiac hypertrophy. A wide range of studies have focused on molecular regulators of cardiomyocyte size, such the calcineurin/NFAT pathway [21]. In contrast, pathways that distinctly control cardiomyocyte shape, such cardiomyocyte elongation often seen after volume overload or endurance exercise, remain underexplored. Here, we show that Nrg1 regulates both size and shape across neonatal rat cardiomyocytes in complementary live-cell and fixed-cell assays, and two lines of hiPSC-derived cardiomyocytes. Using a new metric designed for sensitivity to length and width termed Feret elongation, we identified that in addition to increasing cell area, Nrg1 is a potent inducer of cardiomyocyte elongation.

To identify pathways associated with cardiomyocyte size and shape, we performed proteomic screens in response to hypertrophic ligands ET1, AngII, Nrg1, and IGF1. Partial Least Squares Regression modeling revealed patterns of protein expression and phosphorylation that are co-varied with gene regulation and cell morphology. Based on the strong association of cardiomyocyte elongation with MEK, PI3K, and p38, we performed an additional ligand-inhibitor screen which identified distinct roles of these kinases on myocyte morphology and gene expression. These predictions guided subsequent perturbation experiments with more sensitive fixed-cell high-content microscopy, which identified that Nrg1 mediated PI3K-dependent cell elongation while both PI3K and p38 contribute to cell area. Integrating these data into a logic-based network model demonstrated sufficient mechanisms by which sustained PI3K drives cell elongation, while negative feedback by dual-specificity phosphatases (DUSPs) [22], [23] attenuates p38-driven regulation of cell area. Together, these results highlight how distinct signaling cascades differentially regulate cardiomyocyte size versus shape.

Our proteomic-phenotypic analysis aligns with prior studies. For example, we found that X4E.BP1(T37/T46), Src(Y527), and YAP(S127) were predictive of cell area, consistent with previous studies of cardiomyocyte hypertrophy [16], [17], [18]. In contrast to cell area, only a few pathways have been shown to drive cardiomyocyte elongation (e.g. CT1/LIF, MEK5, β_IV_-spectrin, endurance exercise) or protect against it (CITED4, ERK1/2) [11], [24], [25], [26], [4], [27], [28], [29], [30], [31]. Here by live-cell imaging [13] and fixed-cell immunofluorescence, we find that Nrg1 most substantially increased elongation. Notably, in neurons, Nrg1 is well characterized to promote neurite outgrowth and axonal elongation [32], underscoring its broader role regulating cell shape. Together, these findings decouple “size” from “shape” in cardiomyocytes: PI3K drives both, whereas p38 selectively regulates area.

Although Nrg1-mediated elongation was consistently observed across neonatal rat cardiomyocytes, hiPSC-CMs, and hiPSC-CM^2^ cells, these studies do not fully capture the adult myocardial niche. *In vivo* studies are required to determine the extent to which these cell phenotypes and proteomic signatures drive physiological vs. pathological eccentric cardiac hypertrophy. Still, our data demonstrates that Nrg1 induces several responses associated with physiological hypertrophy including PI3K/Akt activation [7], CITED4 expression [33], [13], and enhanced sarcomere organization associated with increased maturity (**Supplementary Figure 3**) [7]. Further, we found Nrg1 downregulates pathological markers C/EBPβ, CTGF, and Bax [13]. Indeed, a number of studies in rodents and humans have shown that Nrg1 mediates beneficial aspects of endurance exercise, which induce a physiological eccentric hypertrophy [28], [34], [35], [36].

Nrg1 has previously been implicated in cardiac development and disease [37], [38]. In zebrafish, Nrg1 overexpression enhances cardiomyocyte proliferation and supports trabeculation—an essential morphogenetic process required for myocardial growth prior to coronary vessel formation. Disruption of trabeculation results in embryonic lethality or adult-onset dilated cardiomyopathy [39], [40]. In mammals, Nrg1-induced hypertrophy also may be accompanied by some degree of cardiomyocyte proliferation [40], [41], [42], [42], [43], [44]. In humans, chronic exercise has been associated with elevated Nrg1 and physiological hypertrophy [36]. Circulating Nrg1 also rises with heart-failure severity, consistent with a compensatory response to stress [45]. Recombinant human Nrg1 (Neucardin) has completed phase III trials as a treatment for chronic heart failure [46], although those results are not yet available. Phase II trials showed structural and functional improvements [47]. However, not all findings are favorable, with some evidence that Nrg1-induced hypertrophy and ejection fraction may be accompanied by reduced cardiac output [12]. Together, these observations underscore a context- and time-dependent role for Nrg1: acutely adaptive, but potentially maladaptive with chronic exposure.

These findings of distinct regulation of size and shape signaling are currently limited to an *in vitro* context and ligand or pharmacologic perturbations, which motivate future studies. For example, do other more pathologically-associated stimuli of myocyte elongation (LIF, MEK5 [24], [48]) induce distinct pathway mechanisms from Nrg1, which appears more physiological? We perturbed PI3K and p38 with widely-used and validated inhibitors LY944002 and SB203580 [49], [50], but they also have documented off-targets that warrant additional inhibitors and genetic perturbations in future studies. While we tested neonatal cardiomyocytes and two human iPSC-cardiomyocyte lines, it is of interest to expand this study to adult cardiomyocytes and subsequently to in vivo animal models. Future studies should test Nrg1 and downstream pathways implicated here in vivo, incorporating longitudinal analyses to distinguish adaptive from maladaptive remodeling. Examination of Nrg1-induced cell proteomic-phenotype signatures and mechanisms should be tested in endurance exercise [34], which involves physiological eccentric hypertrophy and Nrg1 secretion but incompletely characterized mechanisms. Integration of single-cell transcriptomics or spatial proteomics with machine learning image analysis may help identify how cell size and shape are coordinated at the tissue level.

In summary, this study identifies Nrg1 as a potent regulator of both cardiomyocyte elongation and area, mediated through distinct signaling pathways. PI3K signaling contributes to both elongation and cell area expansion, whereas p38 selectively drives cell area growth. These findings reveal that cardiomyocyte size and shape can be uncoupled mechanistically, advancing our understanding of how hypertrophic remodeling is orchestrated. Leveraging systems-level approaches to selectively modulate maladaptive elongation while preserving adaptive growth may open new therapeutic opportunities for the treatment of cardiac hypertrophy.

## Methods

### Cell Culture

All procedures were performed in accordance with the Guide for the Care and Use of Laboratory Animals published by the US National Institutes of Health and approved by the University of Virginia Institutional Animal Care and Use Committee. Neonatal rat cardiomyocytes were isolated from 1 to 2-day old Sprague Dawley rats using the Neomyts isolation kit (Cellutron, Baltimore, MD). Myocytes were cultured on 96-well Corning CellBIND plates coated with SureCoat (a combination of collagen and laminin, Cellutron) and in plating media (Dulbecco’s modified Eagle media, 17% M199, 10% horse serum, 5% fetal bovine serum, 100 U/mL penicillin, and 50 mg/mL streptomycin).

Human iPSC-derived cardiomyocytes (iCell hiPSC-CMs and hiPSC-CM^2^ from FujiFilm) were cultered on 96-well Corning CellBIND plates (10,000 cells/well) coated with SureCoat and in iCell plating media (FujiFilm). After 48 h of incubation with daily media change (iCell Maintenance Media, FujiFilm), the medium was replaced with Williams E Medium (Life Technologies, A1217601) supplemented with cocktail B supplement (Life Technologies, CM4000) serum-starved for at least 4 h before treatment.

Treatment of ligands and inhibitors were performed in comparison to equimolar DMSO negative controls. Ligands were 10 nM ET1, 1 µM Ang II, 10 nM IGF1, or either 10 or 100 ng/mL Nrg1. Figure 1 and 3 used 10 ng/mL Nrg1 R&D #396-HB, (in vitro EC50 0.06-0.3 ng/mL), motivated by our previous concentration response [13]. However, Nrg1 #396-HB became unavailable from the vendor, so **Figure 4**, **Figure 5, Supplementary Figures 2 and 3** used 100 ng/mL Nrg R&D #377-hB (in vitro EC50 2.5-12.5 ng/mL) as noted in their figure legends. Inhibition of PI3K was performed using 10 μM LY294002, which inhibits multiple isoforms of PI3K as well as off-targets DNA-PK and CK2 with similar ∼1 µM IC50. Inhibition of p38 was performed with 10 μM SB203580, which inhibits p38 at IC50 ∼0.5 µM, which is 100-500 greater potency than other targets LCK and GSK3β. Inhibition of MEK was performed using 0.1 µM PD0325901, which inhibits MEK1/MEK2 with IC50 0.33 nM.

#### Live-cell imaging and PCR

Myocytes isolated and plated as above, with a density of 100,000 myocytes/well. Two days after isolation, myocytes were transfected with GFP under a cardiac myocyte specific troponin T promoter using Lipofectamine 2000 (Invitrogen). Two days after transfection, 5x5 mosaic images of myocytes were collected using automated imaging scripts as described previously [13]. After imaging, myocytes were rinsed and cultured in serum free media containing the indicated ligands and/or inhibitors. 48 hours after stimulation with hypertrophic agonists and inhibitors, follow-up images were recorded and total RNA was purified from the myocytes using the RNeasy Mini kit (Qiagen). Complementary DNA was synthesized from 150 ng of total RNA using the iScript cDNA synthesis kit (Bio-Rad). mRNA lavels of nine genes (Bcl-2, Bax, C/EBPβ, CITED4, VEGF, BNP, IκB, CTGF, and GAPDH) were measured using qPCR (Bio-Rad CFX Connect) using iTaq Universal SYBR Green Supermix (Bio-Rad), 3.75 ng of cDNA, and 400 nM of each primer set. Data were analyzed using the comparative CT method with efficiency correction and measurements were taken from three independent myocyte isolations.

#### Immunofluorescent and high-content imaging

Myocytes isolated, plated (at 30,000 myocytes/well), and cultured as above. To prepare for immunofluorescent imaging, cardiomyocytes were first fixed with 4% paraformaldehyde for 20 min and then permeabilized with 0.1% Triton-X for 15 min. Cardiomyocytes were blocked with 1% bovine serum albumin in PBS for 1 hr, then treated with mouse anti-α-actinin primary antibody (Sigma-Aldrich Cat#A7811, RRID:AB_476766) at a concentration of 1:200 overnight. Cardiomyocytes were blocked with 5% goat serum in PBS for 1 hr, then Alexa Fluor-568- conjugated goat anti-mouse secondary antibody (Thermo Fisher Scientific Cat#A11031, RRID:AB_144696) at a concentration of 1:200 was applied for 1 hr. The cells were stained with DAPI prior to imaging.

High-content imaging was performed on the stained cardiomyocytes using an Operetta CLS High Content Analysis System. These images were processed using CellProfiler using a cellular segmentation algorithm developed previously and validated to within 5% of two independent manual segmentations [14], [51]. Median cell area was used as a representative measure of the cell population in each well, and cells with undetectable cytoplasm were not counted.

#### Single-cell morphology features

To quantify cardiomyocyte shape features, single-cell morphological analysis was performed on CellProfiler-segmented images. Myocytes were filtered based on integrated intensity and percent of neighboring cells touching to ensure inclusion of only healthy, well-isolated cells for accurate shape quantification. For each retained cell, 15 morphological metrics were measured, including 10 size metrics and 5 shape metrics. From two size metrics, a new shape metric was derived:

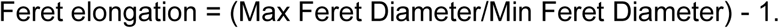

This metric (Feret Elongation) is based on the geometric feature Feret Diameter [19]. Feret Elongation captures the degree of myocyte elongation at all angles, where a value of 0 corresponds to a perfectly non-elongated cell, and higher values indicate greater elongation. Feret elongation is well suited to quantify myocyte elongation because it is scale-independent, rotation-independent, and sensitive to differences between maximum length and minimum width. Scale independence means that the metric remains unchanged if a cell increases or decreases in size proportionally. Additionally, subtracting 1 from the Feret ratio normalizes cells to a baseline minimum width of 1, allowing for comparisons independent of size.

#### Reverse phase protein arrays

Reverse Phase Protein Array (RPPA) analysis was performed at the MD Anderson Cancer Center Functional Proteomics Core to measure the phosphorylation state and changes in protein levels of 172 different proteins. RPPA is a high-throughput method in which antibodies are used to probe spotted arrays of protein lysates and internal controls [52], using antibodies previously validated based by Western blots that exhibit protein bands of correct molecular weight and with positive and negative controls [53].

Neonatal rat ventricular myocytes were cultured in 24-well plates (500,000 myocytes/well). Four days after isolation, myocytes were rinsed and cultured in serum-free media containing 10 ng/mL Nrg1 (R&D #396-HB), 10 nM ET1, 1 µM Ang II, 10 nM IGF1, or serum (10% horse serum and 5% fetal bovine serum). Myocyte protein was isolated at two time points: 1 hour and 48 hours following administration of the ligands according to the protocol on the MD Anderson Cancer Center RPPA Core Facility website. Protein concentration was quantified using the Pierce 660 nM Protein Assay Kit (Thermo Scientific). Cell lysates were submitted to the MD Anderson Cancer Center for analysis. Data was collected from 2 independent myocyte isolations per condition, and all data points were normalized relative to serum-free condition for protein loading.

#### Partial Least Square Regression (PLSR) modeling

Partial Least Squares Regression (PLSR) analysis was performed using MATLAB to infer relationships and visualize covariation among the RPPA panel, phenotypic outcomes, and ligand/timepoint identity within a shared latent space, following its established use for hypothesis generation in high-dimensional systems-level proteomic datasets. The predictor block consisted of 172 proteins across 4 ligands, and the response block comprised 15 phenotypic outputs (11 mRNA transcripts and 4 morphology metrics) across the same ligands. Prior to constructing the model, both RPPA and phenotype data were log-transformed and standardized by computing z-scores across ligands to ensure comparability and eliminate scale differences. PLSR was implemented using MATLAB’s plsregress function. Given the modest number of independent experimental conditions (n=8: four ligands × two timepoints, each with two biological replicates), model validity was assessed by leave-one-condition-out cross-validation, in which both replicates of a given ligand-timepoint condition were withheld together to avoid information leakage between correlated replicates. The number of latent variables was selected by maximizing cross-validated Q² rather than fixed a priori. Statistical significance of model performance was assessed by permutation testing (1,000 iterations of response-label shuffling). Applied to the full 172-protein panel, this procedure did not yield significant predictive value (Q² < 0 across all phenotypes tested; permutation p > 0.05), reflecting the high predictor-to-observation ratio.

All MATLAB scripts used for data preprocessing, model fitting, cross-validation, and visualization are available on GitHub at https://github.com/saucermanlab/neuregulin.

#### Network model construction

A knowledge-based computational model of the distinct cardiomyocyte size and shape signaling network was developed from known direct molecular interaction from the literature, RPPA measurements, phenotypic assays, and PLSR predictions. We employed the logic-based differential equation (LDE) approach [54] to build a predictive framework to explore cardiomyocyte cell area and elongation regulation. A normalized Hill function modeled activation of each node by its upstream reactions. We modeled pathways crosstalk by continuous gates representing “OR” and “AND” logic [54]. The OR gates were used for reactions that modify the node regardless of others and the AND gates for reactions affecting each other. The default reaction parameters in the model are reaction weight (W = 1), half-maximal effective concentration (EC_50_ = 0.5), and Hill coefficient (n = 1.223). The time constant (τ =0.1), initial activation (Yinit = 0), and maximal activation (Ymax = 1) regulate the dynamics of signaling nodes. In conditions with Nrg1 treatment, its input reaction weight was increased (0.02 → 0.3). Further, time constants (tau parameters) were manually calibrated to fit the dynamics of species activation in the experimental data (Table S1). The Netflux software (available on GitHub at https://github.com/saucermanlab/Netflux) has been used to generate a system of LDEs.

#### Statistics

All data are presented as the mean ± SEM. Experiments had a hierarchical design of multiple wells (the experimental unit, randomly assigned to treatment conditions) for each cell isolation (from a neonatal litter composed of 12-16 pups) [55]. Thus, we used linear mixed models [56] accounting for this design with wells as fixed effects and isolations as random effects, with the number of wells and isolations provided in each figure legend. For single-cell measurements, individual cell measurements for each condition are provided in Supplementary Table 3. For all analyses, a P-value < 0.05 was considered statistically significant. For all analyses, a *P*-value < 0.05 was considered statistically significant. Statistical outputs from linear mixed effects models are shown in the Supplementary Text.

The network model, reverse phase protein array, and microscopy data are deposited in FigShare at publication.

## Acknowledgments

We thank Dr. Marieke Jones (Staff Biostatistician, University of Virginia School of Medicine) for statistical consultation and Dr. Alexander Clark (University of Virginia, Department of Biomedical Engineering) for feedback on the manuscript.

## Abbreviations

Akt: protein kinase B;
AngII: angiotensin II;
ANOVA: analysis of variance;
BNP: brain natriuretic peptide;
CITED4: Cbp/p300-interacting transactivator 4;
CM: cardiomyocyte;
CTGF: connective tissue growth factor;
ERK: extracellular related kinase;
ET1: endothelin 1;
GSK3: glycogen synthase kinase 3;
IGF1: insulin-like growth factor;
iPSC: inducible pluripotent stem cell;
MAPK: mitogen activated protein kinase;
MEK: NRCM, Nrg1, neuregulin-1;
p38: PI3K, PLSR, partial least squares regression;
RPPA: reverse phase protein array.

---

**Supplementary Figure 1.**
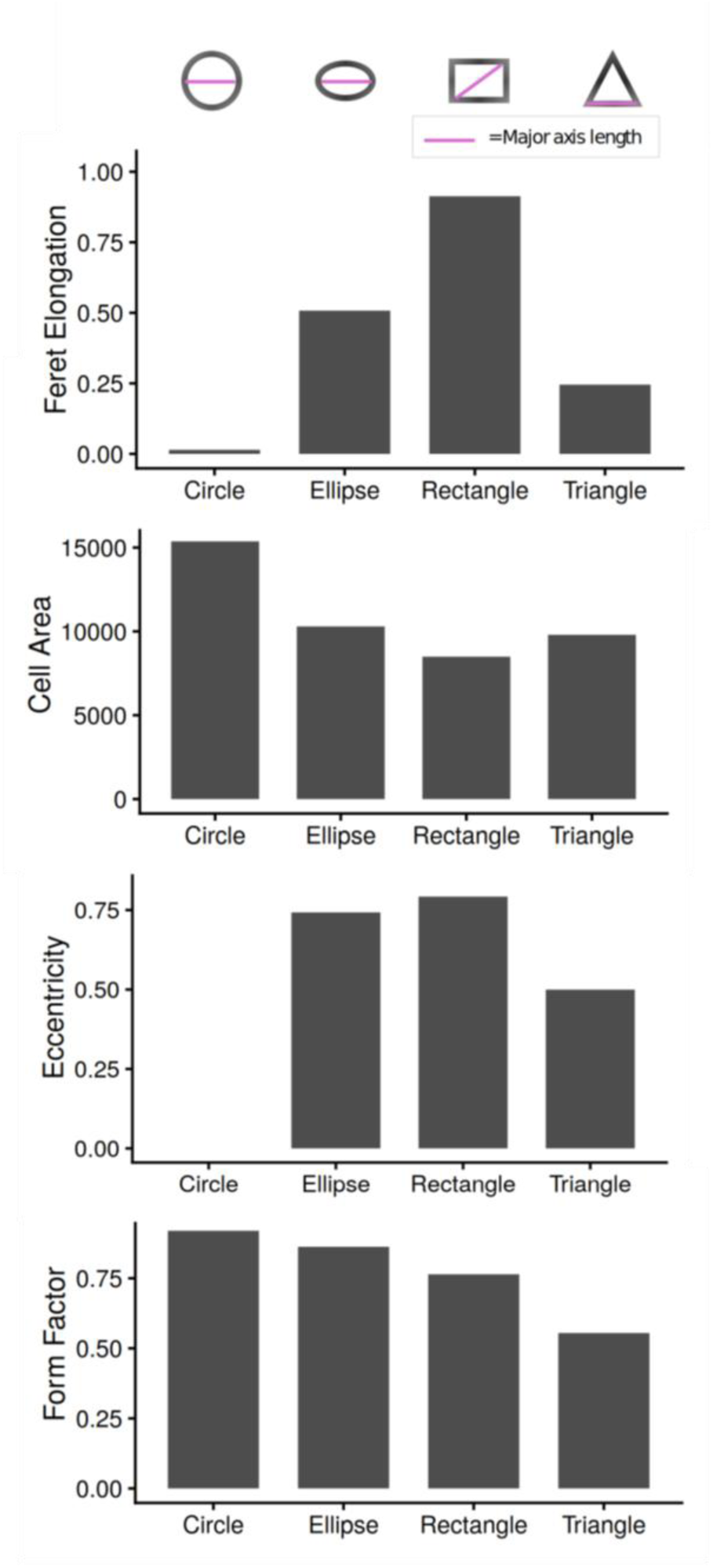
Quantifications using cell morphological features in idealized shapes; circle, ellipse, rectangle, and triangle.

**Supplementary Figure 2.**
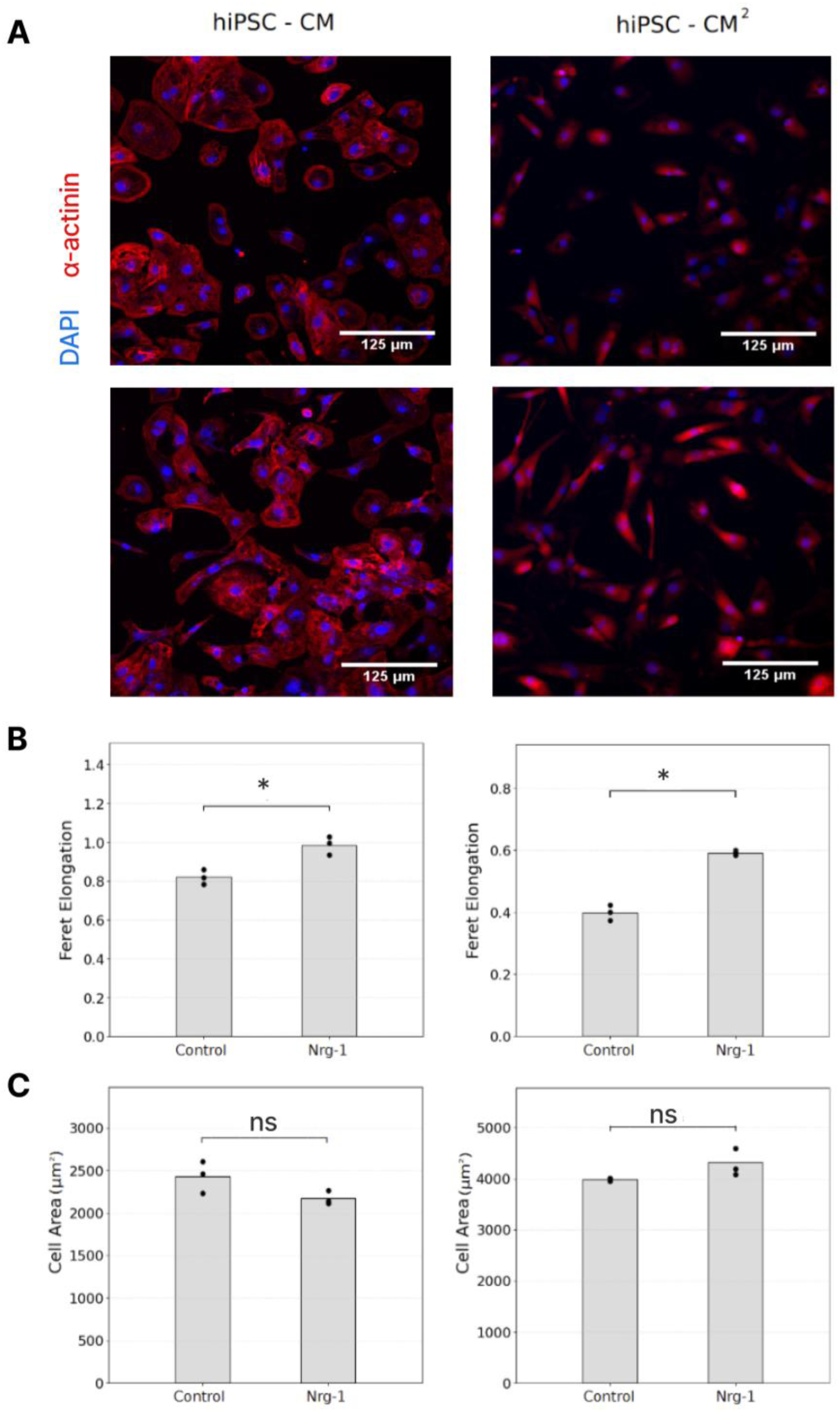
Neuregulin-1 induces elongation of human iPSC-derived cardiomyocytes. (A) Representative images of hiPSC iCell cardiomyocytes and hiPSC iCell Cardiomyocytes² at 48 h after treatment with control or 100ng/mL Neuregulin (Nrg1, R&D #377- HB). Cardiomyocyte marker (α-actinin) and nuclei (DAPI) are shown. (B) Change in Feret Elongation in response to Nrg treatment in hiPSC-CM, and hiPSC-CM². (C) Change in cell area in response to Nrg treatment in hiPSC-CM, and hiPSC-CM². Each point (dot or triangle) reflects the mean of all cells per image, and the bar reflects mean across images. Asterisks denote p < 0.05 from Student’s t-tests. “ns” denotes not significant.

**Supplementary Figure 3.**
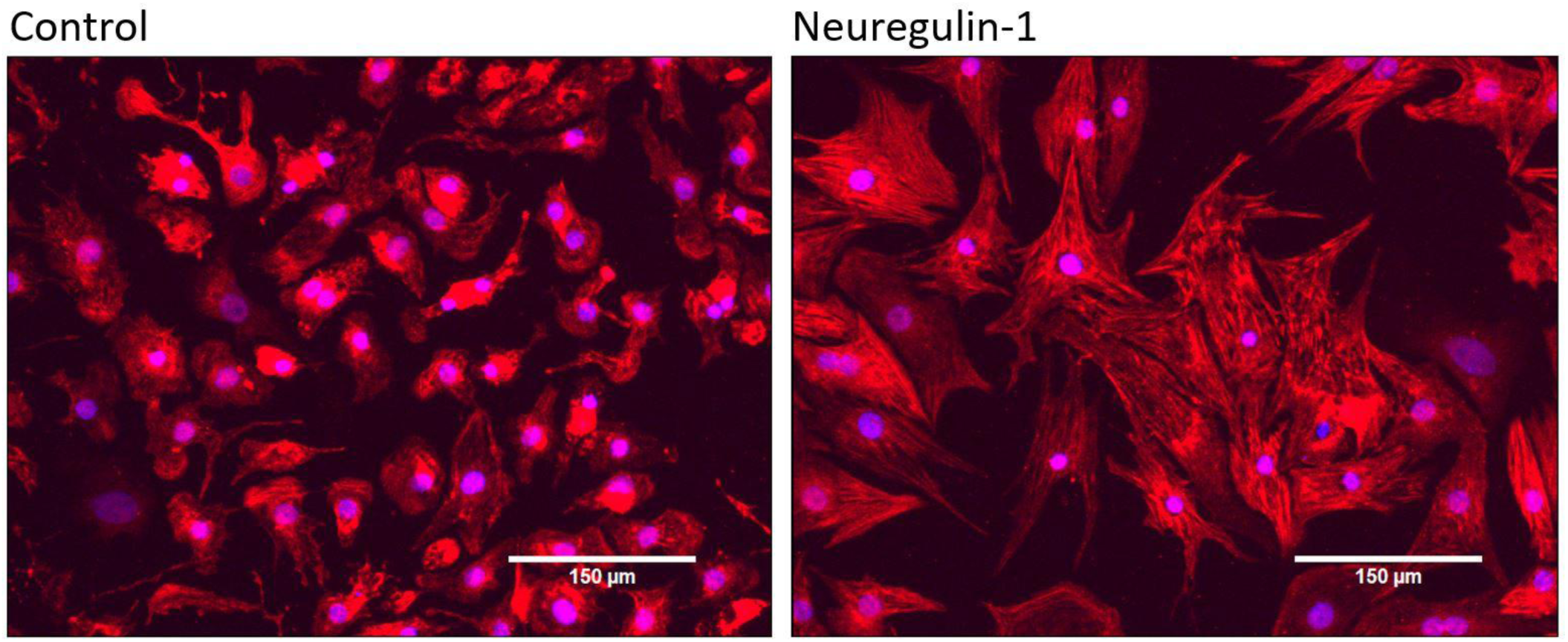
Nrg1-treated neonatal rat cardiomyocytes exhibited formation of organized sarcomeres with visible myofibrils. 100 ng/mL Nrg1 (R&D #377-HB).

**Supplementary Figure 4.**
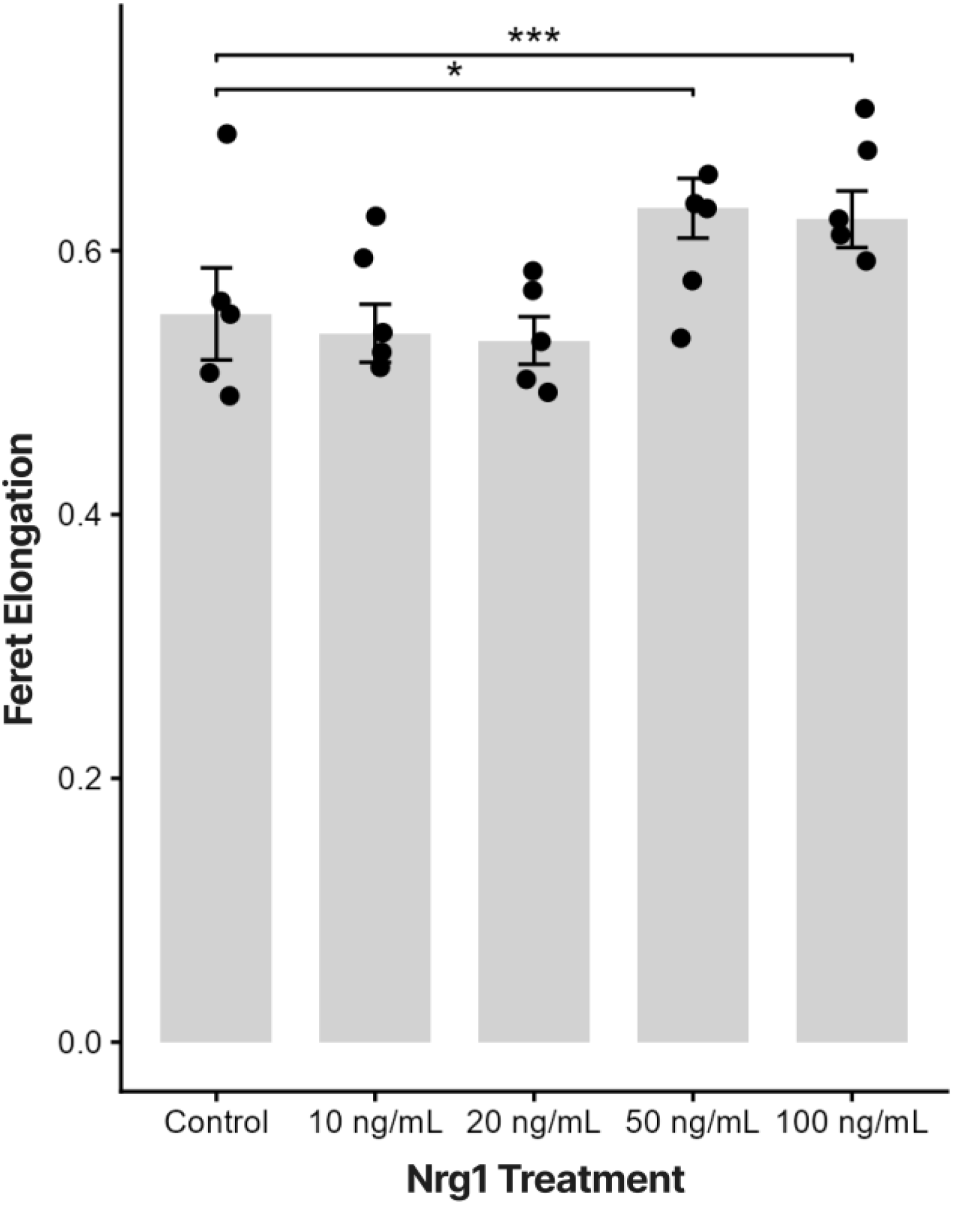
Dose-dependent effect of Nrg1 treatment on cardiomyocyte morphology. Bar graph showing Feret elongation of cardiomyocytes treated with increasing concentrations of Nrg1 (0, 10, 20, 50, and 100 ng/mL). Bars represent mean ± SEM; individual data points are overlaid. *p < 0.05, ***p < 0.001 (linear mixed model, comparing each dose to control; n = 5 per group).

